# The dilemma of underestimating freshwater biodiversity: morphological and molecular approaches

**DOI:** 10.1101/2024.01.23.576836

**Authors:** Alexandra Schoenle, Dominik Scepanski, Alexander Floß, Pascal Büchel, Ann-Kathrin Koblitz, Anja Scherwaß, Hartmut Arndt, Ann-Marie Waldvogel

**Affiliations:** Ecological Genomics, Department of Biology, Institute of Zoology, Biocenter Cologne, University of Cologne, Cologne, Germany; General Ecology, Department of Biology, Institute of Zoology, Biocenter Cologne, University of Cologne, Cologne, Germany

**Keywords:** metabarcoding, morphotypes, species richness, V9 region

## Abstract

**Background:** The Lower Rhine ecosystem has been extensively shaped by human activities, destroying, modifying, and even creating novel water bodies as habitats. Freshwater systems are hotspots of biodiversity and highly complex ecosystems. However, knowledge and description of its biodiversity across all trophic levels is still incomplete and the complexity of interactions remains unresolved, especially below the micro scale. This is even true for important international inland waterways such as the River Rhine. We here document the biodiversity of the Lower Rhine and associated water bodies, spanning from the level of protists up to the level of larger invertebrate predators and herbivores organized in faunal size classes (nano-, micro, meio- and macrofauna). This study is part of a long-term ecological research project (LTER-D REES). Our study includes two riprap sections of the river’s main channel, two oxbows and two gravel-pit lakes in the river’s flood plain. Diversity was assessed through morphotype richness and metabarcoding, as well as accounting for sediment composition.

**Results:** We found high discrepancies between amplicon sequencing variants (ASVs) and morphotype richness in all size classes, as well as a problematic limitation of reference sequences from freshwater organisms in public databases. Irrespective of the size class, we observed habitat specific zoobenthos communities in each of the three investigated habitat types, with few taxa or taxonomic groups overlapping.

**Conclusions:** Our results demonstrate the importance to integrate different methodologies and extend freshwater sequencing efforts for the assessment of biodiversity across all trophic levels, as particularly relevant for long term projects.

## 1. Background

Although freshwaters only account for approximately 0.8 % of the Earth’s surface, yet its habitats sustain around 10 % of all described living species including us humans [1]. In the anthropogenically dominated cultural landscape of Central Europe, the past decade’s intensive utilization of freshwater habitats has been causing degradation, pollution, water extraction, food-web disturbance and invasive species introduction into freshwater habitats, causing a recent biodiversity decline far greater than in most terrestrial ecosystems [1, 2]. Flowing waters and the associated limnic ecosystems of floodplains are particularly impacted by the biodiversity decline and other aspects of global change [3]. As an international stream, the River Rhine is the largest federal waterway in Germany and of corresponding socio-economic importance. Here, clusters of rocks, so called ripraps, can often be found as a result of river straightening [4–6]. The demand of earth-bound resources, of which gravel is particularly available in the grounds of flood plains and thus mined, has furthermore resulted in the creation of artificial lake systems along the river’s course. Despite the imminent threat to freshwater ecosystems and their socio-ecological importance, investments in both research and conservation of freshwater biodiversity lag far behind those in the terrestrial and marine realms [7]. Freshwater systems are hotspots of biodiversity encompassing the genes, populations, species, communities that are relevant for a complex interplay of ecological processes. Knowledge of the species diversity and richness is essential for generating a better understanding of freshwater ecological processes which in turn provide crucial ecosystem services fundamental for human livelihoods and well-being [1]. Numerous studies in different frameworks and habitats have already demonstrated the complex interactions between organisms within the different trophic levels, as well as between size classes [8, 9]. While top-predators like fish are often known to regulate the population of smaller size classes, protists are shown as the essential group for the transfer of energy by connecting lower and higher trophic levels, consequently affecting the whole food web [8, 10]. Many freshwater studies focus on one or few specific taxonomic group [4, 11, 12], but investigations that integrate across all size classes are scarce [8, 13].

Biodiversity monitoring is under development, shifting from a traditional morphological to a genetic based approach for the registration of species. Here, metabarcoding is used as massively-parallelized, high-throughput identification of entire assemblages within an ecosystem [14, 15] and is a non-invasive, cost efficient and rapid tool to monitor biodiversity. However, the inability to differentiate between life stages, intact and living organisms, ingested, or extraneous tissue, to calculate abundances and biomass, or to exclude taxa through unfitted barcodes are remaining obstacles with metabarcoding [14, 16–18]. While enabling quantitative assessments of a community, morphological methods are time-consuming because sampling techniques vary in suitability depending on the different taxa and size classes [19–21]. Thus, a combination of multiple techniques is needed, which subsequently increases the study’s costs and time. Furthermore, correct taxon determination based on morphological characteristics requires a notable amount of knowledge, it is necessary to include at least one expert for each size class to sufficiently cover the community’s diversity.

Given the diversity and richness of species in freshwater systems, biodiversity assessments at habitat-scale and for the magnitude of different freshwater ecosystems is unrealistic to be fulfilled in projects of reasonable size and duration when classical morphology-based identification is considered exclusively. Consequently, the scientific community strives to develop new approaches for a holistic but scalable biodiversity assessment. Here, environmental DNA (eDNA) metabarcoding approaches have been identified amongst the most promising approaches to overcome previous limitations of solely morphological approaches, by their ability to detect an almost complete proportion of the species diversity in a given environmental sample [22–24]. Not only do they have the potential to significantly decrease time, costs and required knowledge but eDNA sequencing rather detects too many than too few taxa inhabiting an area [19–25]. For ecological long-term studies it is particularly relevant to rely on standardized procedures, which are as less error prone as possible, time and cost efficient and independent from exclusive expert knowledge.

The aim of this study was to investigate species diversity from protists up to larger invertebrates organized in size classes (nano-, micro, meio- and macrofauna) from a long-term ecological research site located in Germany at the Lower Rhine in North Rhine-Westphalia including two riprap river Rhine sections, two oxbows as well as two gravel-pit lakes in the floodplain. The aim of the LTER-D project REES is to investigate the eco-evolutionary dynamics along a trophic cascade, integrating species representatives with regards to demography and molecular evolution. Within this study the biodiversity of the targeted REES habitats was initially assessed to obtain a status quo of the water bodies and to create a biodiversity inventory that allows for the choice of representative species along trophic levels for the future development of the long-term study. Diversity was assessed through morphotype richness and metabarcoding, as well as accounting for sediment composition. Furthermore, this study gives insight into how much time, expertise and what kind of mismatch can be expected by targeting biodiversity from a morphological and molecular perspective when people of different expertise and career level are contributing to the assessment.

## 2. Material and Methods

### 2.1 Sampling site

The study area lies within the LTER-D REES site (DEIMS-ID: https://deims.org/554de3a9-1ad9-46e9-9b70-f6e25a799876) at the Lower Rhine in North Rhine-Westphalia (district Rees) in Germany, which resembles the original flood plain area of the River Rhine (partly separated by dikes) including several gravel pit lakes, Rhine oxbows and abandoned meanders, as well as the main river. Littoral benthic samples of four standing water bodies within this area (oxbow Bienen, oxbow Grietherort, gravel-pit lake Reeser Meer Norderweiterung, gravel-pit lake Reeser Meer Süd) and thin epilithic biofilm communities of two ripraps in the River Rhine (Grietherort and Cologne) were collected between 8th of July and 1st of September 2021 (Fig. 1).

**Table 1.** List of the six sampling sites including sampling dates and coordinates.

| **Habitat** | **Site name (Abbr.)** | **Site** | **Sampling date** | **Coordinates (Lat./Lon.)** |
| --- | --- | --- | --- | --- |
| Oxbow | Bienener Altrhein (BAR), Lower Rhine region, Germany | A | 2021-07-08 | 51°48'21.9''N / 6°21'44.9''E |
| Oxbow | Grietherorter Altrhein (GAR), Lower Rhine region, Germany | B | 2021-08-05 | 51°47'29.6''N / 6°19'51.6''E |
| Gravel-pit lake | Reeser Meer Norderweiterung (RMNE), Lower Rhine region, Germany | C | 2021-07-19 | 51°45'41.9''N / 6°26'15.7''E |
| Gravel-pit lake | Reeser Meer Süd (RMS), Lower Rhine region, Germany | D | 2021-07-21 | 51°45'17.8''N / 6°27'32.9''E |
| River Rhine | Riprap Grietherort (RRG), Lower Rhine region, Germany | E | 2021-08-26 | 51°47'25.3''N / 6°19'44.9''E |
| River Rhine | Riprap Cologne (RRC), Lower Rhine region, Germany | F | 2021-09-01 | 50°54'24.0''N / 6°58'43.4''E |

**Fig. 1.**
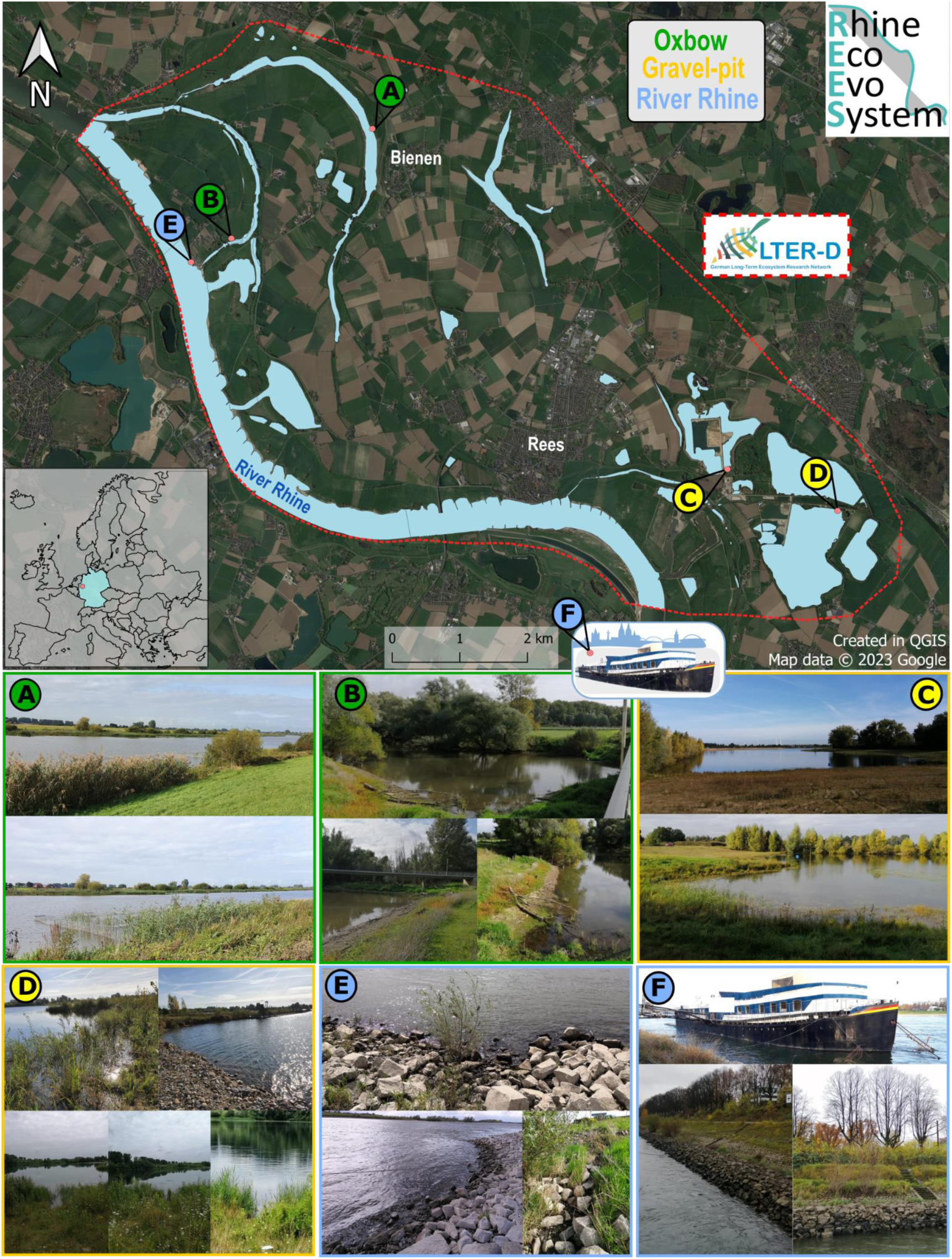
Map and pictures of the six sampling sites (A-F) as part of the LTER-site REES. This site is located at the Lower Rhine in North Rhine-Westphalia (dashed red line, except for site F). The map inlet shows the location of the sampling site in Germany (light blue) within Europe. Samples were taken between June and September 2021. Sampling sites include the oxbows (green) Bienen (A) and Grietherort (B), the gravel-pit lakes (yellow) Reeser Meer Norderweiterung (C) and Reeser Meer Süd (D) and the River Rhine main channel at (blue) Grietherort (E) and Cologne (F). Map was created with QGis (2022) by Tobias Nickel.

While Oxbow Bienen (site A) is an old oxbow only connected to the River Rhine at extreme flood events, oxbow Grietherort (site B) is regularly connected to the main river. Both areas are under nature conservancy, whereby oxbow Bienen (site A) is particularly distinguished as a bird sanctuary. Both site’s shores showed occasional trees and grasses, accumulations of water lilies (*Nymphaea*), common reed (*Phragmites*) as well as nearby stinging nettles (especially site B). Additionally, site A’s vegetation contained further patches of reedmace (*Typha*), mint (*Mentha*), as well as duckweed (*Lemna*) in and around the oxbows shore.

The two **riprap sites** were dominated by basalt boulders ranging from approximately ten to 45 cm in diameter. The riprap in Cologne (site F) and the riprap in Grietherort (site E) were located at the impact slope of the River Rhine. Besides thick biofilm layers overgrowing each submerged rock, no notable macrophytes were observed below water level.

Regarding the two groundwater-fed **gravel-pit lakes,** Reeser Meer Norderweiterung (site C) and Reeser Meer Süd (site D), the habitats’ shores contained more soft sediment than big boulder, while site D showed several rock accumulations above and below the water level. Both lakes are still being used for gravel extraction. With several kilometers distance to the River Rhine and separation by the main dike, both sites’ structural features are unaffected by flooding events; however, the connection via groundwater changes in correspondence with a changing water level of the River Rhine, too.

Both sites shared aquatic patches of pondweed (*Potamogeton*) as well as stoneworts (*Chara*). Both, together with the waterweed *Elodea* were especially dominant at site D’s shore region. While several patches of *Phragmites* grew near site D’s shore only few individuals were observed at the site of gravel-pit lake C. A unique characteristic of site C is the presence of a dense population of the great crested newt (*Triturus cristatus*) and the presence of only one fish species, the sunbleak (*Leucaspius delineatus*), which was first observed in 2019. Before that, the Reeser Meer Norderweiterung had been fish free.

### 2.2 Sampling procedure

At each sampling site, nanofauna (2-20 µm), microfauna (20-200 µm), meiofauna (200 µm-2 mm), and macrofauna (< 2 mm) [26] samples were taken in addition to eDNA samples for metabarcoding analysis.

Macrofauna of the lentic water bodies was sampled in triplicates using a benthos net (mesh size 500 µm) covering 5 m² (oxbow site A) or 0.53 m³ (oxbow site B, gravel-pit lakes site C and D) per replicate. For the other three size classes, sampling corer (diameter 4.4 cm) were used to collect 3.7 cm deep sediment cores together with 7 cm (3.7 cm for site A, Oxbow Bienen due to the low water level) of overlaying water. Ten cores per triplicate (final volume 1000 ml) were filtered through a 500 µm net to remove large particles, leaves and macrofauna. The mixture of one replicate was evenly transferred into four 250 ml plastic beakers (subsamples). For metabarcoding studies, one subsample (250 ml) was filtered through cellulose nitrate filters with a pore size of 0.45 µm (Sartorius Stedim Biotech, Sartorius AG, Göttingen, Germany). Filters were immediately fixated with the salt solution DESS [27] in 50 ml tubes. The other three subsamples (250 ml each) were individually filtered through a meiofaunal sieve (pore size 44 µm), while collecting the filtered water for diversity analyses of protists belonging to the nano- and microfauna. The retained meiofauna on the sieves was suspended in filtered water and transferred into 50 ml tubes. One subsample was fixated with formaldehyde for qualitative analysis, one was fixed in DESS for molecular analyses, and the third was left unfixed for quantitative analyses in the laboratory.

In case of the riprap samples, the benthic community of the submerged rocks was brushed off and sucked in by a pond vacuum cleaner (PONDOVAC 4, OASE GmbH, Hörstel, Germany). To sample eDNA as well as size-classes smaller than macrofauna, biofilms were collected along a transect of defined length with a total surface area of 0.017 m^2^. The collected suspension (~1 L) was processed identical as the previously described sediment samples. For the macrofauna, larger amounts of biofilm were sampled to increase the possibility to collect the entire benthic community. Organisms were retained by a 500 µm net placed over the vacuum’s exit pipe and the volume of run-through water was determined. Sampling continued until approximately 30 L to 40 L were collected. The sampled surface area was measured.

### 2.3 Grain size measurement

Two sediment samples (cores) of the oxbows and gravel-pit lakes were taken to investigate sediment quality of the upper 3.7 cm. Organic matter like leaves or branches were discarded and the wet weight of the sediment was measured. Each sample was dried overnight in a compartment dryer at 60 °C (Memmert GmbH + Co. KG, Schwabach, Germany) and the total dry weight was measured. Different sieves (63 µm, 125 µm, 250 µm, 0.5 mm, 1 mm, 2 mm) were used to determine the relative contribution of each size fraction to the total dry weight. By subtracting the weight of these size fractions from the total dry weight, the amount of fine sediment (< 63 µm) was calculated and the sediment fractions associated to the sediment types gravel (> 2 mm), very coarse (2 mm – 1 mm), coarse (1 mm – 0.5 mm), medium (500 µm – 250 µm), fine (250 µm – 125 µm) and very fine sized sand (125 µm – 63 µm), as well as silt together with clay (< 63 µm) [28].

### 2.4 Morphological identification

Morphological identification of each site’s zoobenthos community was conducted using both light microscopes (Zeiss Axio Lab.A1, Carl Zeiss, Germany) and binoculars (Leica LED2500 stand (Leica Microsystems GmbH, Germany) with stereomicroscope Zeiss Stemi 2000-C attachment (Carl Zeiss AG, Oberkochen, Germany) and attached camera systems (Sony® camera HDR-XR200VE and HDR-XR160E, Sony® Group Corporation, Tokyo, Japan). Specific identification literatures were used for nanofauna [29–31], microfauna [32–38], meiofauna [39–44] and macrofauna [45–54]. The diversity was assessed based on morphotype-richness using determinations of the lowest possible taxon based on literature. If multiple morphotypes were related to one shared taxon, variants were distinguished and numbered to compensate for misidentification.

### 2.5 Metabarcoding

DESS preserved sediment samples were vortexed for two minutes and centrifuged (4000 × g for 20 min at 4 °C, Megafuge 2.0 R, Heraeus Instruments). Environmental DNA was then extracted from 1 g sediment of each replicate sample (a total of 3 g per site) using the DNeasy PowerLyzer PowerSoil DNA isolation kit (Qiagen, Hilden, Germany) according to the manufacturer’s protocol. Prior to the application of the kit, sediment samples were pre-washed with three washing solutions to improve the success of DNA amplification by PCR [55]. Total DNA was quantified using Quantus™ Fluorometer (Promega, Wisconsin, USA). PCR amplifications of the hypervariable V9 region of the 18S rDNA gene was performed with 12.5 µl of the 2X VWR Red Taq DNA polymerase Master Mix (VWR, Erlangen, Germany) and the forward/reverse primer-pair 1389F (5′-TTG TAC ACA CCG CCC-3′) and 1510R (5′-CCT TCY GCA GGT TCA CCT AC-3′) [56]. The PCR mixtures (25 µl final volume) contained 30 ng of total DNA template per site (10 ng per replicate) with 0.35 µM final concentration of each primer. PCR amplifications (98 °C for 30 s; 25 cycles of 10 s at 98 °C, 30 s at 57 °C, 30 s at 72 °C; and 72 °C for 10 min) of all samples were carried out with a reduced number of cycles to avoid the formation of chimeras during the plateau phase of the reaction. Amplifications per site were conducted in triplicates in order to smooth the intra-sample variance while obtaining sufficient amounts of amplicons for Illumina sequencing. PCR products were checked on a 2 % agarose gel for amplicon lengths and fragment sizes were determined by comparison with a 50 bp DNA Ladder (Nippon Genetics Europe, Düren, Germany). Amplicons were then pooled and purified using the FastGene Gel/PCR Purification Kit (Nippon Genetics Europe, Düren, Germany). For subsequent quality measures during data analysis for protists, we used an *in vitro* community (“mock community”) comprising DNA of nine different protist cultures belonging to representatives of the main protist supergroups [57]. We used this mock community to correct for the occurrence of non-reliable sequences. Bridge amplification and paired-end (2 × 150 bp) sequencing of the amplified fragments were performed using a NovaSeq 6000 system at the Cologne Center of Genomics (CCG) in Germany.

### 2.6. Raw sequencing data processing and taxonomic assignment

After sequencing, the raw reads were demultiplexed (parameters: *–no-indels*, –*overlap* 8, *e* 0) and barcode and primer sequences (parameters: *–no-indels*, *-m* 30, *e* 0.2) were clipped using cutadapt version 2.8 [58]. Amplicon sequencing variants (ASVs) were generated by using DADA2 version 1.26 [59] implemented in R v.4.3.0 [60]. Filter and trimming parameters were set to *maxEE* = 1, *truncQ* = 11, *truncLen* = (125, 120), and *maxN* = 0 for quality filtering of the reads, followed by training of error models using the *learnErrors* function of DADA2 with default settings. Dereplication of sequences was done with the *derepFastq* function and the inference of ASVs was done with the *dada* function and default settings. Paired reads were merged with a minimum overlap of 12 nucleotides with *mergePairs*. Chimeric sequences were removed using the *removeChimeraDenovo* function. We used the PR^2^ database version 4.14.0 for taxonomic assignment of ASVs via the pairwise alignment function *usearch_global* of VSEARCH v2.18.0 [61]. All unassigned sequences were removed, keeping only ASVs with a pairwise identity of ≥80% to a reference sequence. First, we divided our dataset into a protist and metazoan dataset. Since this study focuses on zooplankton/zoobenthos, we excluded phototrophic protist sequences (determined on the basis of taxonomic assignment) for the protist dataset including Ochrophyta, Haptophyta, Filosa-Chlorarachnea within the Cercozoa as well as the Cryptomonadales and Pyrenomonadales within the Cryptophyta. Moreover, we removed Metazoa, Archaeplastida and Fungi for the protist dataset. As described by Dünn and Arndt (2023), as a last filtering step, we used a minimum threshold of the previously described mock community by calculating the proportion of the lowest read number of an ASV in the mock community data set that could be assigned to the cultured species with a pairwise identity of 100%. ASVs in the sample data sets with a smaller read number than this calculated proportion were discarded. For the metazoan dataset, besides ASVs assigned to Craniata, we removed terrestrial and marine groups from the final ASV dataset as these might be the result of database limitations. The protist dataset was divided into nano- and microfauna and the metazoan dataset into meio- and macrofauna based on size ranges of taxa found in literature.

### 2.7 Community diversity analysis

Statistical analyses were conducted with R v.4.3.0 [60]. Binary-Jaccard distances were used as a measure of beta-diversity by calling the function *vegdist* within the “vegan” package [62]. The Jaccard distance values were then used for the unweighted pair-group method with arithmetic means (UPGMA) cluster analyses (*hclust* function). Results of the cluster analyses were visualized in dendrograms by using “ggplot2” [63]. Non-metric multidimensional scaling (NMDS) was performed with Jaccard distances using “vegan” to show variation of habitats based on morphotype detection. Observed mean morphotype richness per size classes of respective sampling sites were compared amongst each other via Kruskal-Wallis one-way analysis of variance and Dunn’s post hoc analysis with three replicates per size-class and site (n = 72). Venn diagrams [64, 65] were used to visualize the number of shared and unique ASVs and morphotypes between the three habitats and the different size classes. Heatmaps were created by using the package “ampvis2” [66]. For the heatmaps, only ASVs with a 98-100% sequence similarity to deposited sequences in the PR2 database were kept and clustered to a predicted genus level. With regards to the relative abundance of this ASV sequence similarity range, the 35 most abundant genera per size class were shown in the heatmaps.

## 3. Results

### 3.1 Samples and sediments

In total, 12 samples were taken from the six different sampling sites including four size classes. Three replicates were sampled per size class. All samples could successfully be investigated according to the full spectrum of analysis of this study, except for metabarcoding of the oxbow Grietherort (site B) due to PCR failure. Alongside with the biological samples, sediments of each sampling site were analyzed to better define the habitat characteristics on the micro scale via grain size categories.

Four sites (oxbows and gravel-pit lakes) were characterized by fine sediment, oxbow site A and gravel-pit lake site D by gravelly muddy sand, the gravel-pit site C by gravel sand and the oxbow site B as muddy sandy gravel (Additional file 1, Fig. S1).

### 3.2 Total number of morphotypes and ASVs

This study investigates biodiversity from three different perspectives: morphological diversity (determined by morphotypes), molecular diversity (determined by ASVs with 80-100 % sequence similarity to reference sequences) and the diversity on a predicted genus level (determined by ASVs with 98-100% sequence similarity to reference sequences).

Overall, zoobenthos richness summarized across all size classes and stations showed that out of the total of 191 morphotypes (determined to a lower taxon than the supergroup level) the majority belonged to macrofauna (103 morphotypes) and the lowest number of morphotypes was observed within meiofauna (22 morphotypes). For the protist communities, 24 nanofauna and 42 microfauna morphotypes were detected (Additional file 2, Table S1).

Strict bioinformatic quality control led to a final freshwater eukaryotic dataset of 9,946,586 million reads that clustered into 2,168 amplicon sequence variants (ASVs) containing only ASVs with 80-100% similarity to reference sequences (Additional file 1, Table S2 for more details). Of these ASVs the majority belonged to meiofauna (877 ASVs), followed by macrofauna (434 ASVs). For the protist communities, 425 nanofauna and 432 microfauna ASVs were detected (Additional file 1, Table S3).

Richness obtained by metabarcoding was one order of magnitude higher when directly compared with that recovered by morphological methods (2,168 ASVs versus 191 morphotypes). Metabarcoding always recovered a higher richness than morphotype detection within each of the four size classes (nanofauna: ~20 fold, microfauna: ~10 fold, meiofauna: ~40 fold, macrofauna: ~4 fold). When considering only ASVs with 98-100% sequence similarity followed by a taxonomic clustering on a predicted genus level assignment, the difference between ASV richness and morphotype richness was smaller (nanofauna: ~4 fold, microfauna: ~2 fold, meiofauna: ~3 fold, macrofauna: ~1.3 fold).

### 3.3 Habitat specific zoobenthos richness

The NMDS analysis based on morphotype richness showed that all three different habitats clustered separately with a slight overlap between oxbows and gravel-pit lakes (Fig. 2 A). The mean richness considering all size classes within each of the three habitat types was almost the same in the gravel-pit lakes and the River Rhine (42.3±11.5 ind. and 40±4.05 ind., respectively), while the oxbows showed the lowest mean morphotype richness with 32.5±8.6 individuals (Fig. 2 A Violin plot). Overall, zoobenthos richness with regard to morphotypes across all size classes was not significantly different between habitats (Kruskal-Wallis chi-squared = 3.03, df=2, p-value = 0.22).

**Fig. 2.**
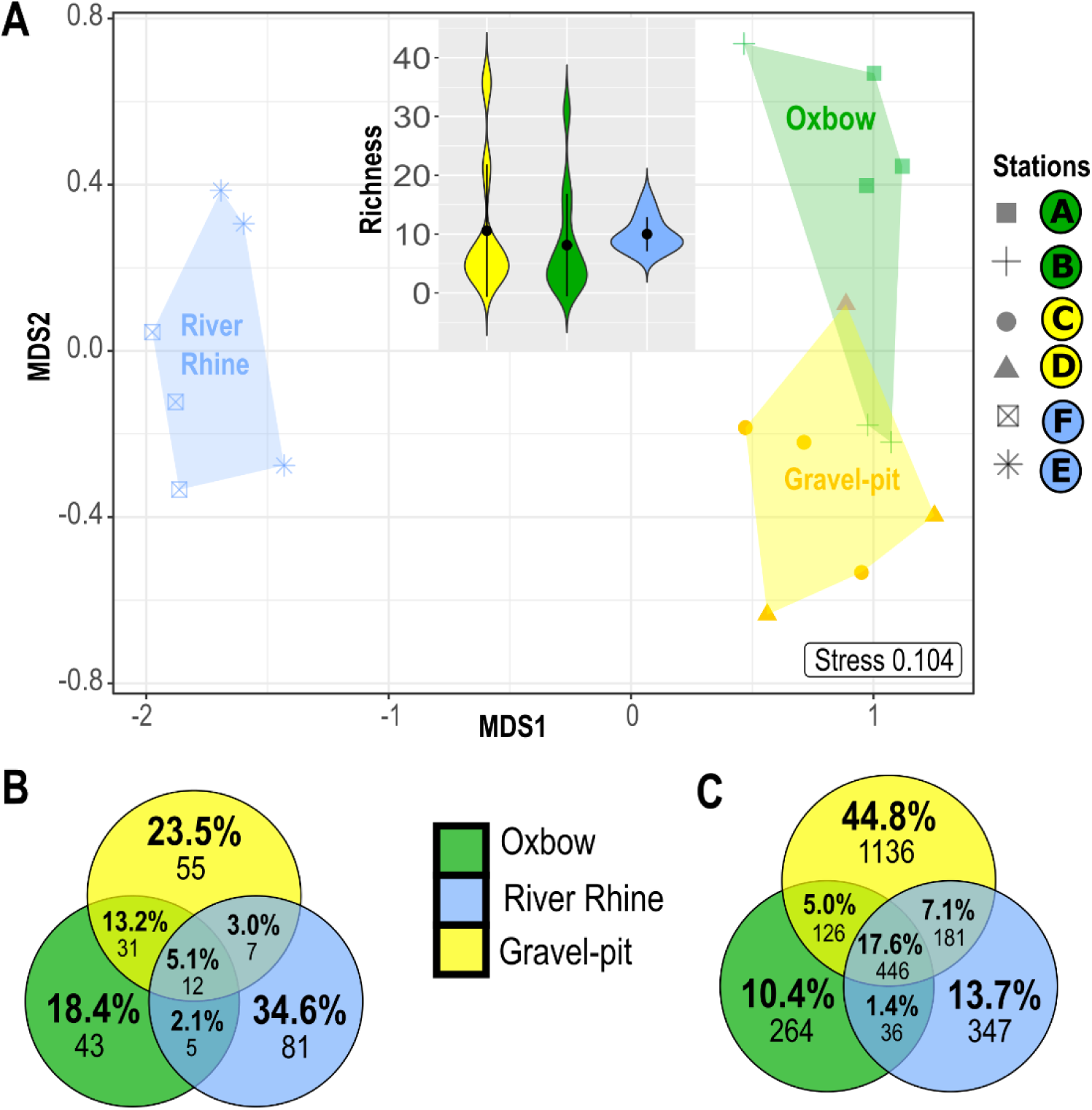
Richness across habitat types. **A.** Non-metric multidimensional scaling (NMDS) analysis of the Jaccard distance matrix computed from morphotypes (n=191 morphotypes) at the six sites (n=3 replicates), associated to three different habitat types (Oxbow, Gravel-pit lakes, River Rhine). The top inset displays the distribution of morphotype richness for the three water body communities. The black dots and bars within violin plots represent means and SDs (replicates per habitat type n=6). **B-C.** Venn diagrams showing the number of unique and shared **(B)** morphotypes and **(C)** ASVs_80-100%_ between the three different habitat types including all four size classes (nano-, micro-, meio- and macrofauna).

The zoobenthic community of the River Rhine was different from all other habitats with 64 unique morphotypes, explaining 34.4 % of the community richness (Fig. 2 B). When comparing morphotype richness of different habitats, it was shown that oxbows inhabit the lowest number of 69 morphotypes, whereas 93 and 86 morphotypes were found in the gravel-pit lakes and the River Rhine, respectively. Only a small proportion (10 morphotypes, 5.4 %) was shared between all habitat types, whereas gravel-pit lakes and oxbows shared 30 morphotypes (16.1%) as indicated in the NMDS plot (Fig. 2 A).

Molecular zoobenthic richness (ASVs_80-100%_) was the highest within the gravel-pit lakes including 904 unique ASVs explaining 41.7 % of the community richness (Fig. 2 C). Comparing ASV_80-100%_ richness between habitats, revealed the oxbows to inhabit the lowest number of 788 ASVs_80-100%_, whereas 1,570 and 912 ASVs_80-100%_ were found in the gravel-pit lakes and the River Rhine, respectively. A higher proportion was shared (404 ASVs_80-100%_, 18.6%) between all habitat types (Fig. 2 C), when compared to morphotype richness (Fig. 2 B). We discovered the overall highest ASVs_80-100%_ richness as well as the highest unique ASVs_80-100%_ richness within each investigated size class (40.6% macrofauna, 45.6% meiofauna, 40.7% microfauna, 35.8% nanofauna) in the gravel-pit lakes (Additional file 1, Fig. S2).

### 3.4 Taxonomic comparison of richness assessed via ASVs and morphotypes

Hierarchical clustering of ASVs_80-100%_ and morphotype richness showed that each habitat formed a separate cluster within the nano-, micro- and macrofauna (Fig. 3 A, C and 4-5 A). For the meiofauna, however, the clustering differed in branching of the gravel-pit lakes, while the cluster for the River Rhine sites was consistent (Fig. 4 A). Overall, nano- and microfauna richness were less similar between sampling sites (Jaccard distances between 0.56-0.93) when compared to the meio- and macrofauna (Jaccard distances between 0.39-0.63) (Fig. 3 A, C and 4-5 A).

**Fig. 3.**
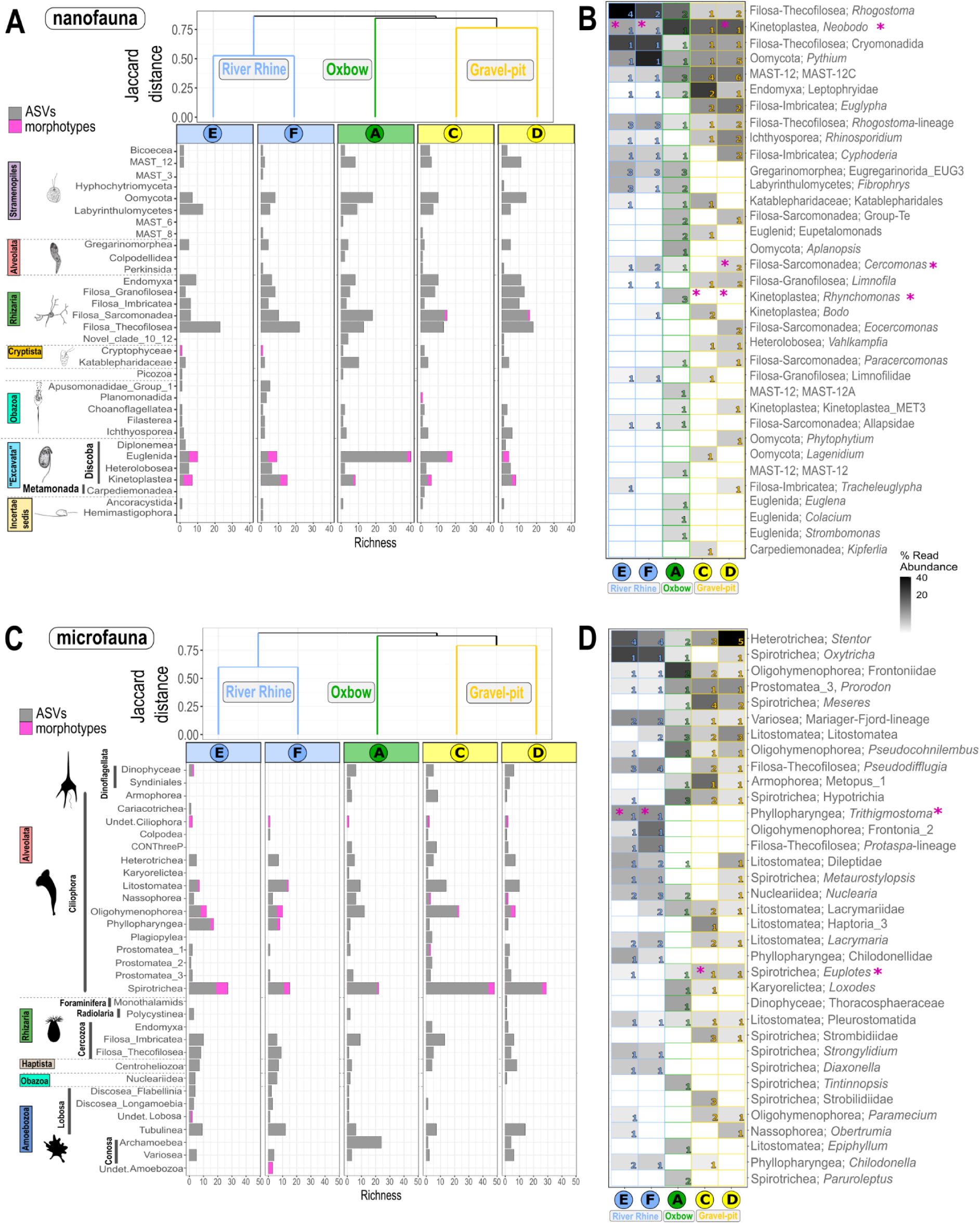
Distribution and community composition of freshwater nanofauna (A-B) and microfauna (C-D). **A and C.** Dendrogram cluster showing the similarity (Jaccard index) of size class communities of the five sediment samples in regard to species richness based on incidence-based data (presence/absence) using UPGMA clustering. Number of freshwater ASV_80-100%_ (grey bars) and number of morphotypes (pink bars) related to the major taxonomic protist groups are shown. Taxonomic groups correspond to class level in the PR^2^ database classification. **B and D.** Heatmap of nano- and microfauna ASVs_98-100%_ with a 98-100% sequence similarity and taxonomic clustering on predicted genus level. Shown are the first 35 most abundant genera (out of 43 nanofauna genera and 79 microfauna genera) with class and genus level. In several cases reference sequences were not assigned to genus level, thus, a higher taxonomic level is shown. Numbers within the heatmap correspond to the number of ASVs assigned to this genus per site. Taxonomic groups correspond to class and genus levels in the PR^2^ database classification. The sequential color code corresponds to the relative abundance of reads with a sequence similarity of 98-100 % assigned to the respective genus to either the nano- or microfauna size class. Pink asterisks indicate, if genera could also be detected morphologically. Nanofauna protist drawings are adapted from literature [30, 67]. Microfauna protist silhouettes are from PhyloPic (https://www.phylopic.org/; T. Michael Keesey, 2023) contributed by Guillaume Dera, 2023 (CC0 1.0 Universal Public Domain Dedication) and Yan Wong, 2013 (Attribution 3.0 Unported, https://creativecommons.org/licenses/by/3.0/).

**Fig. 4.**
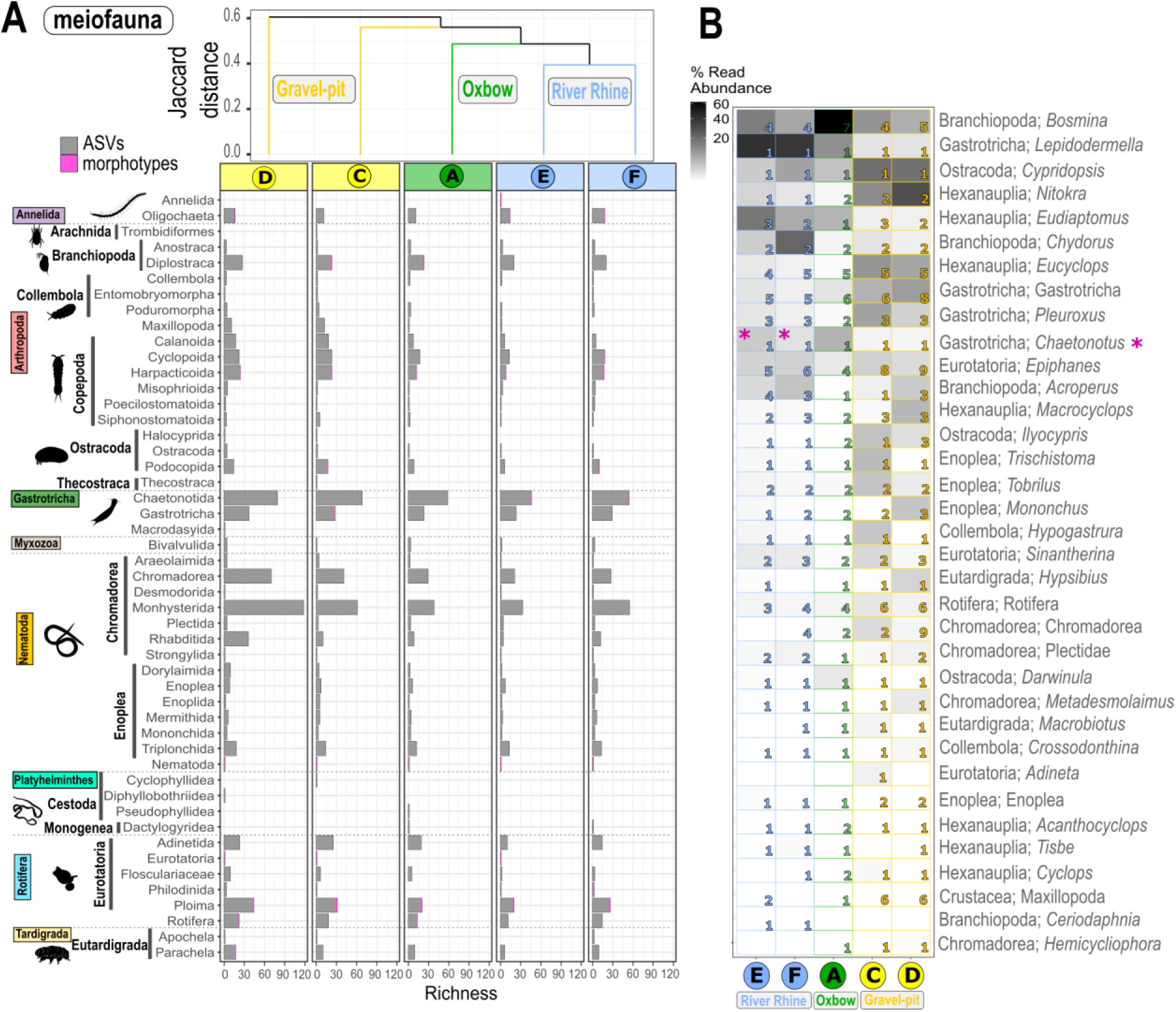
Distributional patterns and community composition of freshwater meiofauna. **(A)** Dendrogram cluster showing the similarity (Jaccard index) of heterotrophic protist communities of the five sediment samples in regard to species richness based on incidence-based data (presence/absence) using UPGMA clustering. Number of freshwater ASVs_80-100%_ (grey bars) and number of morphotypes (pink bars) related to the major taxonomic groups are shown. **(B)** Heatmap of meiofauna ASVs with a 98-100% sequence similarity and taxonomic clustering on predicted genus level. Shown are the first 35 most abundant predicted genera (out of 67 genera) with class and genus level. In several cases reference sequences were not assigned to genus/species level, thus, the deepest taxonomic level such as the order is shown. Numbers within the heatmap correspond to the number of ASVs_98-100%_ assigned to these genera for each of the five sites. Taxonomic genus rank corresponds to genus levels in the PR2 database classification. The sequential color code corresponds to the relative abundance of reads assigned to the genus and is relative to all reads assigned to the meiofauna size classes with a sequence similarity of 98-100 %. Pink asterisks indicate, if genera could also be detected morphologically. Organism silhouettes are from PhyloPic (https://www.phylopic.org/; T. Michael Keesey, 2023). Contributed by Mathilde Cordellier, 2020 (Arachnida), Birgit Lang, 2015 (Collembola), Maxime Dahirel, 2018 (Ostracoda), Michelle Site, 2014 (Nematoda) under License Attribution 3.0 Unported (https://creativecommons.org/licenses/by/3.0/). Contributed by Siel Wellens, 2019 (Copepoda), T. Michael Keesey, 2013 (Branchiopoda), Scott Hartmann, 2013 (Gastrotricha) and Levi Simons, 2023 (Cestoda and Rotifera) under license CC0 1.0 Universal Public Domain Dedication. Contributed by Julie Blommaert, 2020 (Rotatoria) under license Attribution-ShareAlike 3.0 Unported. Contributed by B. Duygu Özpolat, 2016 (Oligochaeta) under license Attribution-NonCommercial-ShareAlike 3.0 Unported.

**Fig. 5.**
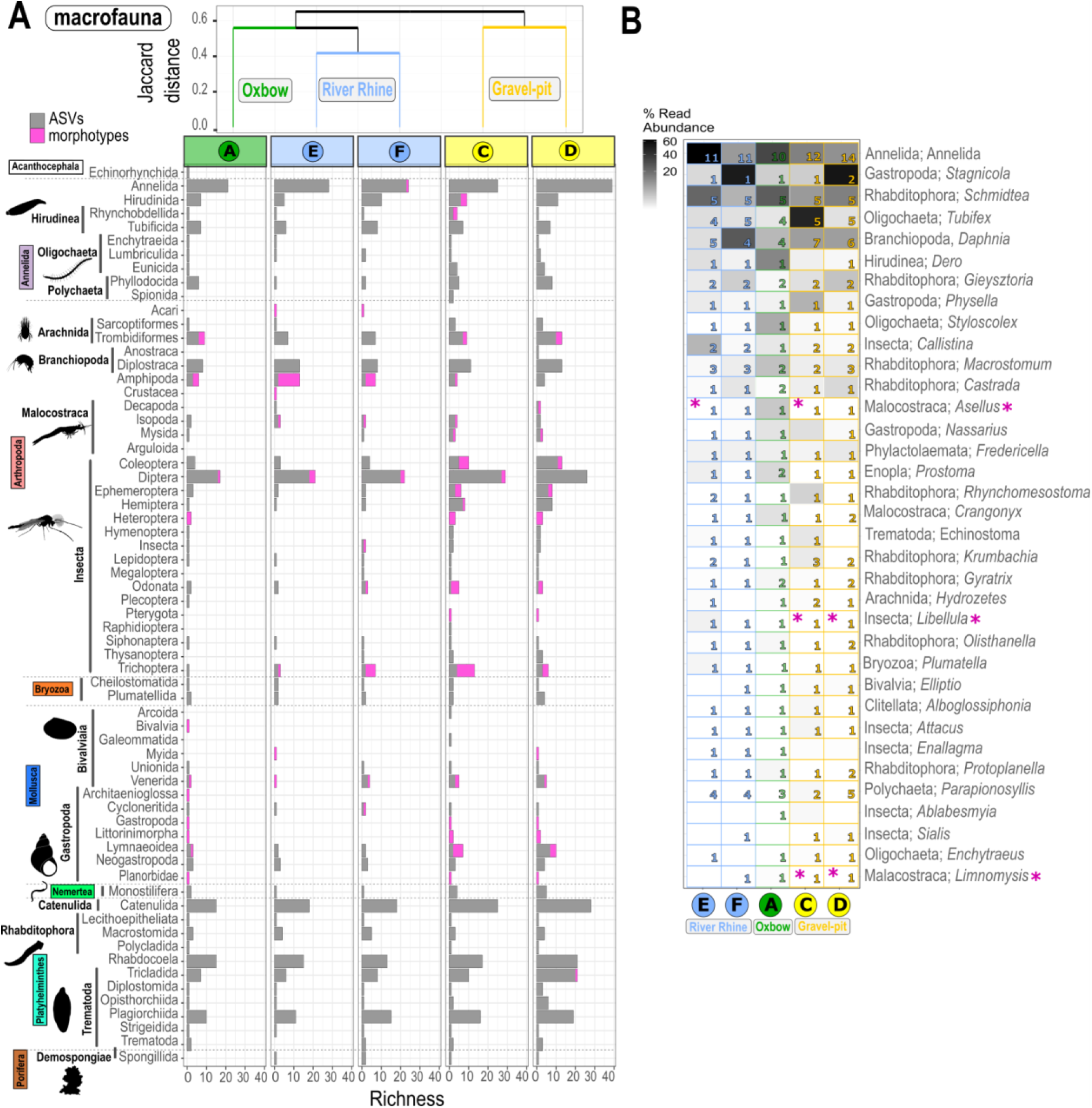
Distributional patterns and community composition of freshwater macrofauna. **(A)** Dendrogram cluster showing the similarity (Jaccard index) of macroinvertebrate communities of the five sediment samples in regard to species richness based on incidence-based data (presence/absence) using UPGMA clustering. Number of freshwater ASVs_80-100%_ (grey bars) and number of morphotypes (pink bars) related to the major taxonomic macroinvertebrate groups are shown. **(B)** Heatmap of macrofauna ASVs with a 98-100% sequence similarity and taxonomic clustering on predicted genus level. Shown are the first 35 most abundant genera (out of 115 genera) with class and genus level. In several cases reference sequences were not assigned to genus/species level, thus, the deepest taxonomic level such as the order is shown. Numbers within the heatmap correspond to the number of ASVs_98-100%_ assigned to these genera for each of the five sites. Taxonomic genus rank corresponds to genus levels in the PR2 database classification. The sequential color code corresponds to the relative abundance of reads assigned to the genus and is relative to all reads assigned to the macrofauna size classes. Pink asterisks behind the taxonomic names indicate, if morphotypes down to genus level could also be identified. Pink asterisks indicate, if genera could also be detected morphologically. Organism silhouettes are from PhyloPic (https://www.phylopic.org/; T. Michael Keesey, 2023). Contributed by Nathan Jay Baker, 2022 (Branchiopoda), Kamil S. Jaron, 2022 (Insecta), Tauana Cunha, 2021 (Gastropoda) and Guillaume Dera, 2023 (Rhabditophora, Trematoda, Porifera) under license CC0 1.0 Universal Public Domain Dedication. Contributed by Denis Lafage, 2020 (Malacostraca), Katie Collins, 2020 (Bivalvia) and Mathilde Cordellier, 2020 (Arachnida) under License Attribution 3.0 Unported (https://creativecommons.org/licenses/by/3.0/). Contributed by B. Duygu Özpolat, 2016 (Hirudinea, Oligochaeta) under license Attribution-NonCommercial-ShareAlike 3.0 Unported (https://creativecommons.org/licenses/by-nc-sa/3.0/).

A direct comparison of ASVs_80-100%_ against morphotypes based on Class level revealed that the morphotype coverage of supergroups was incomplete (Fig. 3 A, C and 4-5 A). After filtering for ASVs with a 98-100% similarity to reference sequences (Additional file 1, Fig. S3 and S4), only 20.5% of the ASVs_98-100%_ were left for the nanofauna (87 ASVs_98-100%_; 432,502 reads; 43 genera, 51 species; Fig. 3 B), 29.4% of the ASVs_98-100%_ for the microfauna (127 ASVs_98-100%_; 523,950 reads; 79 genera, 89 species; Fig. 3 C), 14.8% of the ASVs_98-100%_ for the meiofauna (130 ASVs_98-100%_; 1,421,912 reads; 67 genera, 73 species; Fig. 4 B) and 30.0% of the for the macrofauna community (130 ASVs_98-100%_; 1,841,620 reads; 78 genera, 85 species, Fig. 5 B). The percentage of taxa detected with both approaches ranged from 3.8-44.3%, when only ASVs_98-100%_ with 98-100% sequence similarity were considered. When considering all ASVs_80-100%_ from 80-100% sequence similarity, the percentage of shared taxa was always below 10% with one exception of 14.9% (Additional file 3, Table S4).

#### 3.4.1 Nanofauna

When considering all ASVs with a sequence similarity of 80-100% for the nanofauna community, ASVs_80-100%_ could be assigned to 32 orders. However, only morphotypes belonging to five out of these 32 classes belonging to five supergroups and Incertae sedis could be identified, including morphotypes belonging to the Filosa Sarcomonadea (Rhizaria), Cryptophyceae (Cryptista), Kinetoplastea and Euglenida (“Excavata”), Ancyromonadida (Incertae sedis) (Fig. 3 A). Except for the Ancyromonadida, we also detected a high ASV_80-100%_ richness for these division levels, where morphotypes were found. The highest morphotype richness could be detected for Kinetoplastea and Euglenida, while especially samples from the oxbow revealed a high euglenid ASV_80-100%_ richness (Fig. 3 A). Out of the 24 nanofauna morphotypes, only eight could be determined down to genus/species level, including kinetoplastids (*Neobodo designis*, *Rhynchomonas nasuta*), cercozoans (*Cercomonas* sp.), euglenids (*Petalomonas* sp., *Entosiphon* sp., *Peranema* sp.), cryptophyceans (*Goniomonas* sp.), planomonadids (*Ancyromonas* sp.) (Additional file 2, Table S1). When considering ASVs with a sequence similarity of 98-100% with a taxonomic clustering on predicted genus level, only three genera out of the 35 most abundant ASVs_98-100%_ could also be detected morphologically, however, with great discrepancies when looking at site level matches (e.g. *Rhynchomonas* and *Cercomonas*) (Fig. 3 B). The most abundant nanofauna genus, the cercozoan *Rhogostoma,* based on read abundance, could not be detected morphologically in the samples (Fig. 3 B). The second most abundant genus, the kinetoplastid *Neobodo,* was detected in all habitats and sites by metabarcoding, but morphotype detection revealed this genus to be present only in the River Rhine and one gravel-pit lake (site D).

#### 3.4.2 Microfauna

When considering all ASVs with a sequence similarity of 80-100% for the microfauna community, ASVs_80-100%_ could be assigned to 32 classes belonging to five supergroups. However, only morphotypes belonging to ten out of these 32 classes could be identified, including morphotypes belonging to the Ciliophora (Litostomatea, Nassophorea, Oligohymenophorea, Phyllopharyngea, Prostomatea 1, Spirotrichea), Dinoflagellata (Dinophyceae) and Amoebozoa (Fig. 3 C). Ciliophora had the highest ASV_80-100%_ richness at all sites, with the majority of morphotypes assigned to this group (37 out of 42 total morphotypes). Within the Ciliophora except for the Nassophorea and Prostomatea 1, we also detected a high ASV_80-100%_ richness for these division levels, where morphotypes were found. The highest morphotype richness within the microfauna community could be detected for the Spirotrichea, while also the highest ASV_80-100%_ richness was observed for this group (Fig. 3 A). Only four amoebozoan morphotypes were found, although the number of ASV_80-100%_ and richness was much higher. While we recovered ASV_80-100%_ for several divisions within the rhizarians, we were not able to detect them morphologically. Out of the 42 microfauna morphotypes, 21 could be determined down to genus/species level, including species belonging to the Spirotrichea, Phyllopharyngea, and Oligohymenophorea (Additional file 2, Table S1). When considering ASVs with a sequence similarity of 98-100% with a taxonomic clustering on predicted genus level, only two genera out of the 35 most abundant ASVs_98-100%_ could also be detected morphologically, namely the genera *Trithigmostoma* and *Euplotes* (Fig. 3 D).

#### 3.4.3 Meiofauna

When considering all ASVs with a sequence similarity of 80-100% for the meiofauna community, ASVs_80-100%_ could be assigned to 49 orders. However, only morphotypes belonging to 12 out of these 49 orders could be identified, including morphotypes belonging to the Oligochaeta (Annelida), Diplocostraca (Branchiopoda), Cyclopoida and Harpacticoida (Copepoda), Chaetonotida (Gastrotricha), unidentified nematodes (Nematoda), unidentified Eurotatoria, Ploima and Philodinida (Rotifera), Parachela (Tardigrada) (Fig. 4 A). The highest ASVs_80-100%_ richness was found within the Chaetonotida (Gastrotricha) and Monhysterida (Nematoda). While a few morphotypes could be detected for the Chaetonotida (Gastrotricha), no morphotypes could be detected for the Monhysterida (Nematoda). Out of the 22 meiofauna morphotypes, only eight could be determined down to genus/species level, including Eurotatoria (*Brachionus*, *Euchlanis*, *Philodina*, *Synchaeta*), Diplocostraca (*Alona*, *Eurycercus*), Ostracoda (*Eucypris virens*) and Gastrotricha (*Chaetonotus*) (Additional file 2, Table S1). When considering ASVs with a sequence similarity of 98-100% with a taxonomic clustering on predicted genus level, only one out of the 35 most abundant ASVs_98-100%_ could be detected morphologically, namely *Chaetonotus* sp. (Fig. 4 B).

#### 3.4.4 Macrofauna

When considering all ASVs with a sequence similarity of 80-100% for the macrofauna community, ASVs_80-100%_ could be assigned to 61 orders. However, only morphotypes belonging to 28 out of these 61 orders could be identified, including morphotypes belonging to Annelida (Hirudinea, Rhynchobdellida), Arachnida, Branchiopoda, Malocostraca, Insecta, Gastropoda and Bivalvia (Fig. 5 A). The highest ASVs_80-100%_ richness was found within the Rhabdocoela and Catenulida (Platyhelminthes), undetermined Annelida and Diptera. Except for the Platyhelminthes Rhabdocoela and Catenulida, where no morphotype could be detected, a few morphotypes were found for the other orders with a high ASVs_80-100%_ richness. For Amphipoda, Odonata and Trichoptera the morphotype richness was greater at some sites compared to the ASVs_80-100%_ richness. Out of the 103 macrofauna morphotypes, 51 could be determined down to genus/species level, including Gammaridae (e.g. *Dikerogammarus* sp.), Isopoda (e.g. *Asellus aquaticus*), Mysidae (e.g. *Limnomysis benedeni*), Odonata (*Libellula depressa*), Hemiptera (*Notonecta* sp.), Gastropoda (e.g. *Anisus vortex*, *Bithynia* sp., *Lymnaea stagnalis*) (Additional file 2, Table S1). When considering ASVs with a sequence similarity of 98-100% with a taxonomic clustering on predicted genus level, only three out of the 35 most abundant ASVs_98-100%_ could be detected morphologically, namely *Asellus aquaticus, Libellula depressa and Limnomysis benedeni* (Fig. 5 B).

## Discussion

Several studies have already compared biodiversity assessments via morphological approaches and molecular approaches such as metabarcoding within specific taxonomic groups in different ecosystems [68–71]. The performance and outcomes of biodiversity surveys have been examined with a particular focus on the comparative effectiveness of morphology and DNA-based approaches of specific taxonomic groups highlighting discrepancies that may arise in the application of these methods [72–74]. It has also been shown that the precision and taxonomic resolution of biodiversity surveys can be increased by eDNA surveys [71, 75, 76]. Within our study, the percentage of shared taxa between both approaches eDNA (ASVs_80-100%_) vs. morphology (morphotypes) was always below 10%. Within a study targeting phytoplankton, zooplankton and macroinvertebrates, the number of taxa shared between both approaches was also low with 7-9% [13]. When we considered only ASVs with a sequence similarity of 98-100% for our study, the number of shared taxa reached up to 32%. One example of a perfect match of both applied methods is the phyllopharyngean cyrtophorid ciliate genus *Trithigmostoma,* a common freshwater ciliate, known to feed on diatoms and filamentous microbes. Within the two riprap sites (E and F) high read abundances of the cercozoan species *Rhogostoma minus* were detected by metabarcoding. This delicate thecofilosean could not be identified with the morphological approach during the sampling period, but was identified in the river in earlier studies.

Single primer pairs or combinations of primer pairs sequencing the same or different regions have been used within metabarcoding studies targeting different taxa [77–81]. For protists the 18S rRNA is used as a universal marker to assess protist biodiversity targeting mainly the hypervariable V4 or V9 regions (~420 bp and ~130 bp, respectively) in metabarcoding studies [56, 78, 82, 83]. For several protist taxa the V9 region can be used to detect down to the species level [84], while for others this region was only sufficient to distinguish down to the genus level [85]. The hypervariable V9 region has also been used to investigate the overall metazoan community composition (e.g. Liu and Zhang, 2021). While for a variety of metazoan species the highly conserved mitochondrial region cytochrome oxidase COI is used as a reliable marker gene for identification, several difficulties challenging distinct species identification using COI as the marker gene were addressed by the barcoding community in the past [87–89]. One is the principle of the “barcoding gap”, the lack of a significant difference between intraspecific and interspecific diversity [90]. This phenomenon was also observed for various fish groups which cannot be determined by COI barcoding [91, 92]. Thus, the 12S, 16S and 18S regions are used as reliable marker genes down to species level for fish communities [93–95], while a larger amount of water might be needed to be filtered in order to detect fish eDNA. A reliable taxonomic assignment of nematode species is based on the 18S rRNA together with the COI [96]. The choice of a primer pair for metabarcoding studies is dependent on the research question, the targeted taxonomic group, the sequencing technique (e.g. short vs. long reads) and the available project budget. While investigating different size classes and taxonomic groups within this study, we looked at different levels of molecular diversity, including the molecular diversity based on ASVs with 80-100 % sequence similarity to reference sequences (ASV_80-100%_) and the diversity on a predicted genus level based on ASVs with 98-100 % sequence similarity to reference sequences followed by a taxonomic clustering based on genus level (ASV_98-100%_). These ASVs_98-100 %_ were used for a direct comparison between the morphological diversity based on the morphological approach (morphotypes) and molecular diversity on a predicted genus level based on metabarcoding (ASVs_98-100 %_). While a comparison between ASVs_80-100%_ and morphotypes revealed a distinct mismatch in richness estimates for all size classes, a comparison between morphotypes and genus-clustered ASVs_98-100%_ resulted in the number of morphotypes similar to the number of ASVs_98-100%_. Nevertheless, many major taxonomic groups in all size classes could not be recovered with the morphological approach and on the other hand, many ASVs_98-100%_ of larger organisms have not been found within our morphological detection. Within this study different taxa were most likely recorded under the same morphotype, likely underestimating the true diversity of each site.

Metabarcoding can only be as precise as its database [16, 18, 22]. Not only must a database include enough correctly assigned taxa to potentially find new invasive species, the aspect that many public databases are error-infected makes the availability of well-curated databases an important prerequisite [18]. Several morphotypes identified to species level in this study have been missing in the database. One example is the detected ciliate genus *Aspidisca* for which sequences could be found in the used reference database, but not for the common species identified by the morphological approach. One advantage of metabarcoding is that it is less reliant on taxonomic expertise [14, 69] including many taxa currently not identifiable by expert taxonomists [97–99]. We found *Dikerogammarus* morphotypes only to be present in the samples from the Ripraps in the River Rhine, which are seen are seen as an emerging ecosystem with new substrate-specific combinations and can be dominated by non-native species such as *Dikerogammarus villosus/haemobaphes* [4, 100, 101]. However, metabarcoding only indicated the presence of Gammaridae in the other two habitats (oxbows and gravel-pit lakes), namely *Gammarus pulex* (ASV with 100% sequence similarity) and *Gammarus tigrinus* (ASV with 100% sequence similarity). While no V9 sequence for *Dikerogammarus* was present in the used reference database, reblasting these two *Gammarus* ASVs (read abundance of 394 and 55) at GenBank showed a 100% similarity to many species belonging to the family of Gammaridae, but only a 94.46% similarity to *Dikerogammarus* species. Especially for protists (nano- and microfauna), a combination of molecular and morphological approaches is needed to gain the highest possible community resolution. Another example is the ciliate genus *Stentor,* which is common in the River Rhine at Cologne [11] and was highly abundant with regards to reads from the metabarcoding approach, but could not be detected by our morphological approach at the sampling date. This might be due to the used sampling technique with a vacuum pond cleaner. Besides the ciliate *Stentor*, other biofilm associated taxa like other protists, rotifers and gammarids have been observed within this study by using both approaches, as well as larger sessile organisms, such as the freshwater clam *Corbicula* sp..

Irrespective of the size class, we observed habitat specific zoobenthos communities in each of the three investigated habitat types, with few taxa or taxonomic groups overlapping. The gravel-pit lakes showed the highest overall molecular diversity (ASV_80-100%_) of zoobenthos, while the overall morphological diversity was similar for the gravel-pit lakes and the River Rhine. Although ripraps are considered to provide low physical complexity on a broader scale if used as the environment’s dominant structure, we found the highest unique morphotype richness in the ripraps of the River Rhine with a distinct community composition when compared to the other habitats of the present study. While studies on riprap communities are still limited, it is known that ripraps create several microhabitats with a diverse fauna on a microscale [102, 103]. Oxbows and gravel-pit lakes provide a different habitat for organisms of the different size classes. While the taxa in the River Rhine are highly impacted by the flow velocity and receives organisms from a vast catchment area, the zoobenthos communities of the other two habitats are e.g. influenced through vegetation and substrate composition [4, 104–108]. Whereas patches of macrophytes in the shore area of lakes are generally known to provide distinct ecological niches and food sources benefitting the zoobenthos diversity [105, 109, 110], the high occurrence of macrophytes (*Potamogeton* sp., *Chara* sp., *Elodea* sp.), especially at gravel-pit lake site C’s (Reeser Meer Norderweiterung) shoreline could be one explanation for the higher ASV_80-100%_ within the gravel-pit lakes when compared to the River Rhine. While meiofauna morphotypes, especially nematodes, were underrepresented due to the needed taxonomic expertise to identify genera or species morphologically, we recovered a high genetic diversity of nematodes belonging to the order Monhysterida and Chromadorea as well as Chaetonotida with our metabarcoding approach (ASV_80-100%_) within the gravel-pit lakes. Only for the macrofauna, the morphotype richness was highest in the gravel-pit lakes, while for the other size classes (i.e. the nano- and microfauna) the highest unique morphotype richness was documented from the River Rhine. Surface-associated taxa, which occur attached to particles, such as flattened ciliates, might have resisted the mixing current and therefore did not go into suspension [111, 112]. Moreover, replicates of the morphotype richness for the River Rhine had a much more constant habitat structure, when compared to the gravel-pit lakes, which might also explain the much higher standard deviation of the morphotype richness between the replicates for the gravel-pit lakes and oxbows compared to the River Rhine. Differences in sediment composition and macrophytes at each site might strengthen the differences in community richness between those two gravel-pit lakes, together with the fact, that at the Reeser Meer Norderweiterung (site C) only the sunbleak *Leucaspius delineatus* was present, while the Reeser Meer Süd (site D) is inhabited by many fish species including the European perch (*Perca fluviatilis),* the common roach (*Rutilus rutilus)* and the northern pike (*Esox lucius)*.

### Perspectives for biodiversity monitoring

Biodiversity assessments based on metabarcoding might not replace the traditional morphological approach until sequence availability in databases has been extended dramatically, but is suggested to be used as a complementary tool for biodiversity monitoring [70, 71, 113, 114]. The combination of morphotype detection and metabarcoding was particularly relevant for the biodiversity assessment of bodonids present in all three investigated habitats (genera *Neobodo*, *Rhynchomonas*, *Dimastigella*). This order of kinetoplastids is known to harbor a very high degree of genetic diversity as compared to whole orders of higher eukaryotes suggesting large numbers of cryptic individual species within bodonids [115].

While traditional morphological approaches as well as metabarcoding both have their advantages and limitations, metagenomics might promise a new omics approach for biodiversity assessment to overcome the limitations of metabarcoding such as primer dependency and PCR amplification biases. While metagenomics does not cover the abundance aspect, automated image processing with AI-driven identification of species and simultaneous recording of their size (biomass) is the only viable solution for long-term biodiversity monitoring and will allow for the first time to quantify interaction networks for entire food webs. The present and further acquisition of reference genomes will help discover the genetic mechanisms underlying organisms’ responses to their natural environment and with this support conservation efforts [116]. Reference genomes from a wide range of species are required to map genomic variability and ultimately contribute to the conservation of genetic diversity. Various international initiatives aim to generate reference genomes representing global diversity [117]. With high quality reference genomes, genomic diversity of individuals from the same species can be cost-efficiently unraveled by resequencing and aligning against it [118].

## Declarations

### Ethics approval and consent to participate

Not applicable.

### Consent for publication

Not applicable.

### Availability of data and materials

The data analyzed in this study are deposited at the Sequence Read Archive SRA (SRP481383) with SRA Accessions SRR27420739-SRR27420743, BioProject ID PRJNA1061115, BioSamples SAMN39255222-SAMN39255226.

Preliminary results have been presented at the ECOP-ISOP conference in Vienna, Austria, in 2023 [119].

### Authors’ contributions

HA, AMW, AS contributed to conception and design of the study. DS, AK, PB, AF, AS, HA were involved in the sampling. PB and AF cultivated and sequenced isolated protist cultures. AK processed the meiofauna samples and DS analyzed the macrofauna samples. AS performed the bioinformatic analysis of morphological qualification and quantification data. AS conducted the lab analyses of the sediment samples (next-generation-sequencing). AS performed the bioinformatic analyses of the NGS data. DS and AS wrote the first draft of the manuscript. All authors contributed to manuscript revision, read, and approved the final manuscript.

## Acknowledgements

We are very grateful to Ulrich Werneke and the team of the NZ Kleve for managing sample registration. We thank Rosita Bieg, Brigitte Gräfe and Anke Pyschny for their valuable technical support. We thank the Company Holemans for the permission to sample the two gravel-pit lakes. We acknowledge the Ecological Research Station Rees of the Institute of Zoology, University of Cologne, that served as major research infrastructure for the field work of this study.

