## Additional file 1 for "The dilemma of underestimating freshwater biodiversity: morphological and molecular approaches"

#### A Supplemental figures

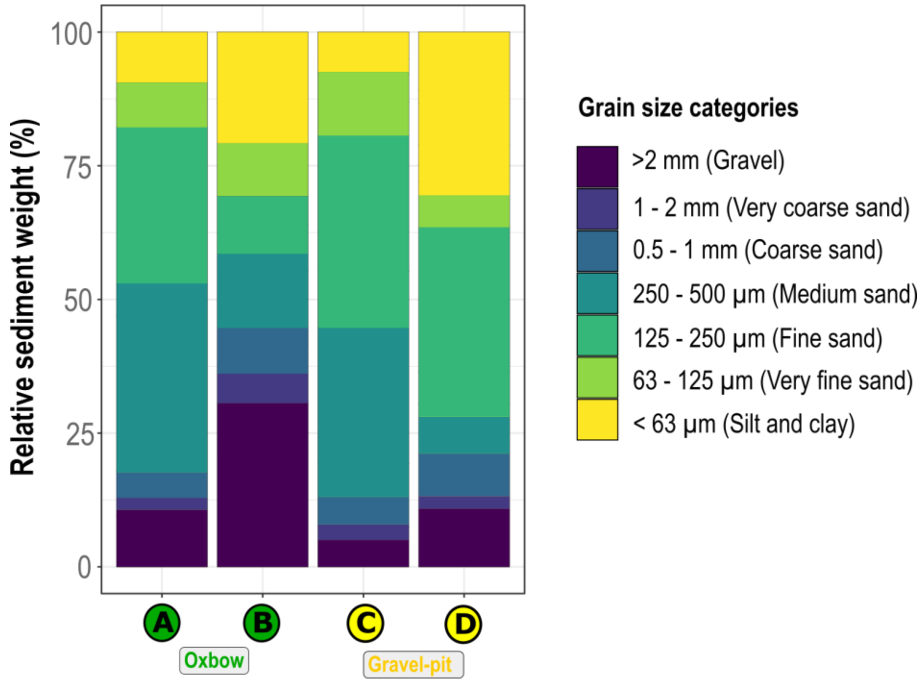


**Figure S1**. **Sediment composition.** Relative sediment weight composition (%) until 3.7 cm depth for the two oxbows (A and B) and gravel-pit lakes (C and D). Sites at the River Rhine (E and F) were not included here, because riprap areas were sampled.

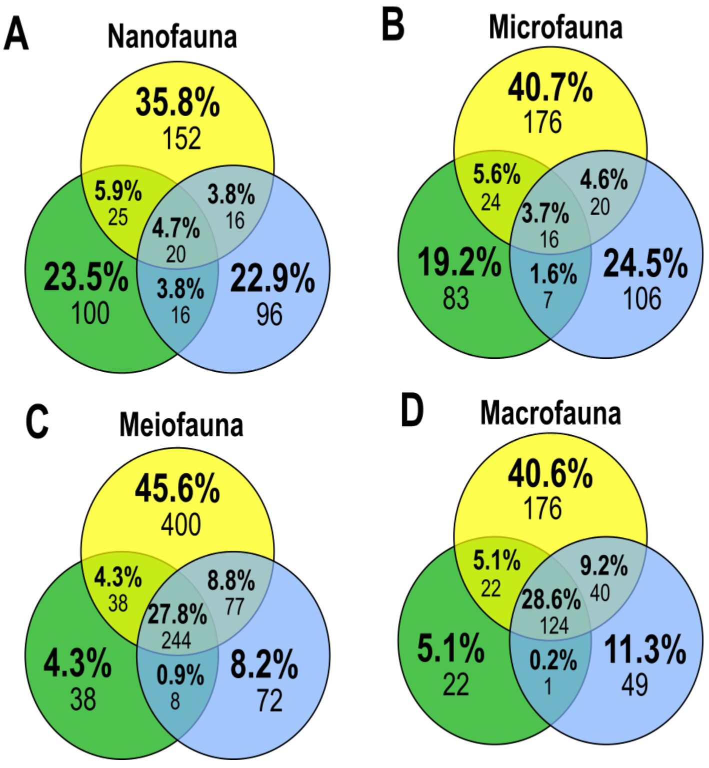


**Figure S2. Shared and unique communities within each size class.** Venn diagrams **(A-D)** showing the number of unique and shared freshwater ASVs$_{80\text{-}100\%}$ between the three different habitat types for all four size classes (**(A)** nano-, **(B)** micro-, **(C)** meio- and **(D)** macrofauna).

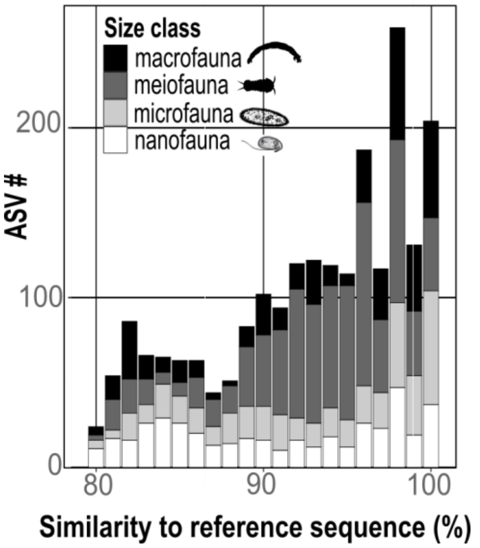


**Figure S3. Similarity to reference sequences.** Similarity plot (80-100%) of eDNA richness to total referenced eukaryotic rRNA diversity in PR$^2$ database per size class. Proportion of ASVs per size class is color coded.

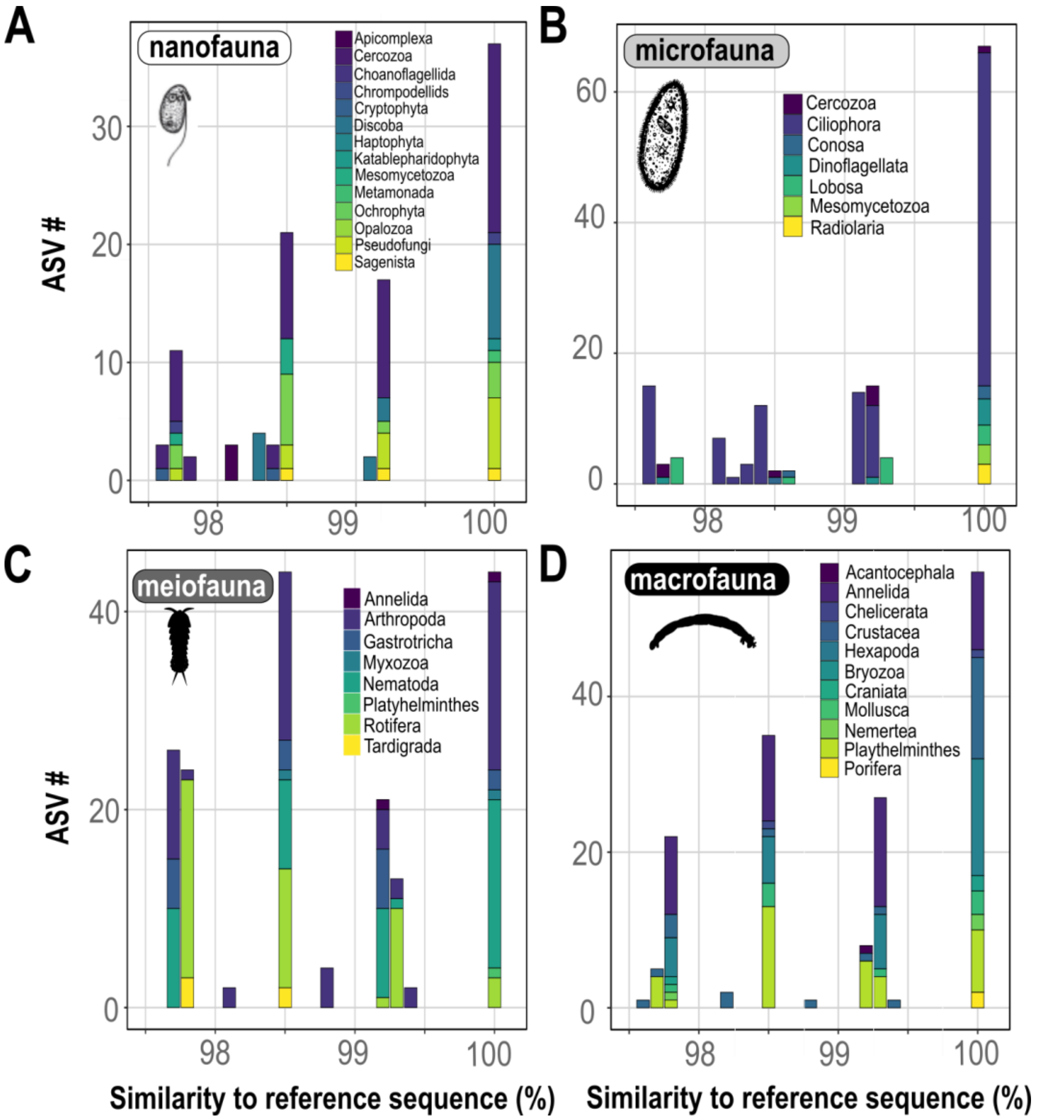


**Figure S4. Similarity to reference sequences within different taxa.** Similarity plot (97.5-100%) of eDNA richness to total referenced eukaryotic rRNA diversity in PR$^2$ database separated into size classes.

### B Supplemental tables

**Additional file 2: Table S1**. **Morphotype richness per size class.** (xlsx file)

**Table S2**. **Metabarcoding filters.**

| **Filter** | **Kept** | **Number of ASVs / Percentage of total** | **Number of reads / Percentage of total** |
|---|---|---|---|
| none | total | 39,199 (100%) | 28,235,163 (100%) |
| no hit | only eukaryotes | 38,831 (99.1%) | 28,229,693 (99.98%) |
| p-identity <80% | p-identity of 80-100 % | 16,882 (43.1%) | 24,701,963 (87.5%) |
| | only freshwater metazoans | 1,311 (3.3%) | 7,468,055 (26.5%) |
| | only protists | 12,229 (31.2%) | 15,378,827 (54.5%) |
| | only heterotrophic protists | 9,866 (25.2%) | 4,171,129 (14.8%) |
| | only heterotrophic protists larger than mock read threshold | 857 (2.2%) | 2,768,575 (9.8%) |

**Table S3**. **Number of reads and ASVs$_{80\text{-}100\%}$ per site and size class (percentage identity 80-100%) after filtering.** Nanofauna with a total of 425 ASVs$_{80\text{-}100\%}$ and 1,281,140 reads, Microfauna with a total of 432 ASVs$_{80\text{-}100\%}$ and 1,197,238 reads, Meiofauna with 877 ASVs$_{80\text{-}100\%}$s and 4,988,078 reads, Macrofauna with 434 ASVs$_{80\text{-}100\%}$ and 2,479,977 reads. Oxbows: A, Gravel-pit lakes: C and D, River Rhine: E and F.

| Site | Size class | No. of reads | No. of ASVs$_{80\text{-}100\%}$ |
|---|---|---|---|
| A | Nanofauna | 10,7361 | 161 |
| C | Nanofauna | 337,559 | 118 |
| D | Nanofauna | 463,334 | 138 |
| E | Nanofauna | 76,189 | 101 |
| F | Nanofauna | 296,850 | 111 |
| A | Microfauna | 82,634 | 130 |
| C | Microfauna | 412,187 | 163 |
| D | Microfauna | 362,525 | 120 |
| E | Microfauna | 79,316 | 118 |
| F | Microfauna | 260,576 | 99 |
| A | Meiofauna | 442,874 | 328 |
| C | Meiofauna | 1,266,201 | 455 |
| D | Meiofauna | 1,764,520 | 641 |
| E | Meiofauna | 449,260 | 281 |
| F | Meiofauna | 1,065,223 | 368 |
| A | Macrofauna | 103,355 | 169 |
| C | Macrofauna | 1,042,947 | 239 |
| D | Macrofauna | 963,017 | 309 |
| E | Macrofauna | 105,874 | 169 |
| F | Macrofauna | 264,784 | 183 |

**Additional file S3: Table S4**. **Summary of richness and shared taxa per size class**. Table contains information on sediment composition, habitat, site, mean morphotype richness, ASV richness (80-100% sequence similarity and 98-100% sequence similarity) as well as the percentage of shared taxa for both ASV richness approaches. (xlsx file)
